# Convergent evolution in *Arabidopsis halleri* and *Arabidopsis arenosa* on calamine metalliferous soils

**DOI:** 10.1101/459362

**Authors:** Veronica Preite, Christian Sailer, Lara Syllwasschy, Sian Bray, Ute Krämer, Levi Yant

## Abstract

It is a plausible hypothesis that parallel adaptation events to the same environmental challenge should result in genetic changes of similar or identical effects, depending on the underlying fitness landscapes. However, systematic testing of this is scarce. Here we examine this hypothesis in two closely related plant species, *Arabidopsis halleri* and *Arabidopsis arenosa*, which co-occur at two calamine metalliferous sites harbouring toxic levels of the heavy metals zinc and cadmium. We conduct individual genome resequencing alongside soil elemental analysis for 64 plants from 8 populations on metalliferous and non-metalliferous soils, and identify genomic footprints of selection and local adaptation. Selective sweep and environmental association analyses indicate a modest degree of gene as well as functional network convergence, whereby the proximal molecular factors mediating this convergence mostly differ between site pairs and species. Notably, we observe repeated selection on identical SNPs in several *A. halleri* genes at two independently colonized metalliferous sites. Our data suggest that species-specific metal handling and other biological features could explain a low degree of convergence between species. The parallel establishment of plant populations on calamine metalliferous soils involves convergent evolution, which will likely be more pervasive across sites purposely chosen for maximal similarity in soil composition.

## 1. Introduction

Most plants cannot rapidly escape hostile environments. Thus, they present powerful models for the study of adaptation. Remarkably, some plant species contain multiple populations that have evolved the ability to thrive in the harshest environments, for example extreme drought, solar radiation, heat, salinity, low nutrient availability and toxic soil concentrations of heavy metal ions. Metalliferous (M) soils are defined as rich in at least one class B and borderline trace metal element [1], are usually nutritionally imbalanced [2, 3], and arise either through geological (e.g. ancient outcrop) or human (e.g. mining, metal smelter) activity. Such soils are generally toxic to plants and host a sparse, species-poor characteristic vegetation of adapted, often endemic extremophiles, so-called metallophytes [4].

Several members of the *Arabidopsis* genus have been described as pseudo-metallophytes, i.e. harbouring populations on both M and nonmetalliferous (NM) soils, namely *Arabidopsis halleri* [5, 6], *A. arenosa* [7-9] and *A. lyrata* [10]. Among these species, only *Arabidopsis halleri* is widespread on calamine-type M soils of Central and Eastern Europe as well as in East Asia and has thus become a model organism for the study of evolutionary adaptation to challenging soils. Calamine soils are defined as containing high levels of zinc (Zn), which are geologically accompanied by the metals cadmium (Cd), lead (Pb) and occasionally copper (Cu). *Arabidopsis halleri* is a diploid (2*n* = 18), stoloniferous perennial and obligate outcrosser with a haploid genome size of approximately 260 Mbp [11]. On both M and NM soils, *A. halleri* exhibits Zn, and regionally also Cd, hyperaccumulation, defined as the ability to accumulate > 3,000 μg Zn g^-1^ dry leaf biomass or > 100 μg Cd g^-1^ dry leaf biomass in its natural habitat [5, 6]. Experimental studies in synthetic hydroponic media have demonstrated species-wide hypertolerance to both metals in comparison to the closely related species *A. lyrata* and *A. thaliana* [12, 13]. These same studies also established that *A. halleri* accessions originating from calamine M soils exhibit enhanced Zn and/or Cd hypertolerance, which is likely the result of local adaptation. Importantly, the basal metal tolerance present in all plants, which enables them to acclimate to local fluctuations in soil composition, does not allow survival on calamine M soils [14].

Generally, *A. arenosa* is absent from most calamine M soils and is known as a so-called metal excluder, i.e. a plant that maintains normal Zn and low Cd concentrations in its above-ground biomass in natural populations [15, 16]. While *A. halleri* and *A. arenosa* generally occupy differing edaphic niches, both species are rarely found together at few calamine M sites [17] in Eastern Europe and at some NM sites (our unpublished observations), suggesting that their populations can occasionally undergo convergent niche shifts. Our information on tolerance to and accumulation of calamine-type metals in *A. arenosa* and its intra-species variation is still sketchy [7, 8, 18, 19].

Work over the past two decades has established a first understanding of the genetic basis of species-wide metal hypertolerance and hyperaccumulation in *A. halleri* in comparison to closely related species [11, 20, 21]. However, no causal genetic locus governing within-species variation in tolerance to calamine-type metals has been identified to date. With this study, we aimed to detect convergent genomic footprints of selection at two calamine M sites, each by comparison to a NM site in their vicinity, in both *A. halleri* and *A. arenosa*. We thus took advantage of a few exceptional cases where both species have adapted to the same sites and thus similarly composed soils. Individual genome resequencing of 64 individual plants from 8 populations, followed by high-density genome scans for selective sweeps, identified a handful of compelling candidate genes under selection at both of the two site pairs or in both of the two species. Notable among these, we identify the *A. halleri Cysteine Protease-Like 1 (CPL1)* locus as a candidate for convergent selection in both population pairs. We show that this gene exhibits a series of convergent derived sequence variants in individuals originating from M sites, and appears to have undergone a loss-of-function in populations at NM sites.

## 2. Methods

### (a) Field sampling, and plant and soil materials

Root-proximal soil samples were collected for multi-element analysis from *Arabidopsis halleri* (L.) O’Kane and Al-Shehbaz ssp. *halleri* and *Arabidopsis arenosa* ssp. *arenosa* (L.) Hayek at four field sites in September 2015 and May 2016 (7 to 10 individuals per species and site; see table S1; see [6] for a description of sites and methods for soil sample collection and processing). Additionally, five to ten leaves per individual were placed in a 2-mL polypropylene tube for later DNA isolation, immediately frozen in liquid nitrogen (MVE vapor shipper, Chart, Minnesota, USA) and stored in liquid nitrogen. For experiments under controlled growth chamber conditions, we collected about 40 L of soil (≤ 0.3 m depth) at the M site Miasteczko Śląskie (Mias; see table S1) in May 2016 (see below). All-purpose greenhouse soil (Minitray, Einheitserde, Sinntal-Altengronau, Germany) was used as NM control soil. Plants were grown from cuttings of *A. halleri* individuals collected at Mias and Zakopane [6] and maintained in the greenhouse (Ruhr-Universität Bochum, Germany, [6]) and from *A. arenosa* seeds collected at the two field sites Mias and Zakopane (Zapa).

### (b) Plant cultivation under growth chamber conditions

Seedlings of *A. arenosa* and vegetative clones of *A. halleri* (see Supplementary methods) were pre-cultivated for 17 days in 1:1 (v/v) peat:sand (round pots, 5 cm Ø, 3.5 cm depth, 50 mL volume) in a climate-controlled growth chamber (20°C/17°C, 10 h light at 100 μmol m^-2^ s^-1^; Grow Banks, Arabidopsis BB-XXL.3, CLF Plant Climatics, Wertingen, Germany). Subsequently, plants were transferred into experimental treatment soils (3 volume parts of field-collected M Mias or MN greenhouse soil, each mixed with 1 volume part of sand; square pots 7 x 7 cm width, 8 cm depth, 300 mL volume) and cultivated in the growth chamber for another 6 weeks. Pots were arranged in trays (separated by species and treatment soil; 16 to 23 pots per tray), and plants were watered with tap water (poured from above) when needed (2 to 3 times per week, preventing waterlogging). Positions and orientation of trays were re-arranged randomly once per week. Photographs were taken at the start, after 3 weeks and at the end of the experimental cultivation period. At harvest, plant survival was scored and fresh above-ground biomass was determined for each plant. The biomass between experimental groups was assessed using a generalized linear mixed effect model with fresh biomass as dependent variable, genotype (site of origin: Mias M site vs. Zapa NM site) and treatment (experimental soil type of exposure: control-soil vs. M soil) as fixed predictors. Individual plants and genotype nested in treatment were set as random factors. The significance of each variable as well as the interaction between genotype and treatment was tested with type II x^2^ based likelihood-ratio tests (based on inverse gaussian distribution with the link function 1/mu^2^; glmer and ANOVA functions in R-packages lme4 [22] and car [23], respectively).

### (c) Analysis of soil samples and DNA extraction

Soil pH, as well as extractable and exchangeable concentrations of Al, B, Ca, Cd, Cr, Cu, Fe, K, Mg, Mn, Ni, P, Pb, S and Zn in soil samples were determined as described [6]. Frozen leaf tissues were lyophilized overnight (Alpha 1-4 LSC plus, Martin Christ LCG, Osterode am Harz, Germany) and subsequently homogenized with a single ceramic bead (3 mm Ø; Precellys Beads, Peqlab, Erlangen, Germany) in a Retsch mixer mill (Type MM300, Retsch, Haan, Germany) for 2 × 1.5 min at 30 Hz. For each individual sample 9 to 15 mg of dry leaf powder was weighed into a 2 mL polypropylene tube and mixed thoroughly with 0.9 mL CTAB buffer, followed by DNA extraction according to [24] with small modifications (see Supplementary methods). DNA quality was verified by spectrophotometry and agarose gel electrophoresis, and DNA was quantified using the dsDNA HS assay (Q32854) following the manufacturer’s instructions with an incubation time of 20 min (Qubit 3.0, ThermoFisher Scientific, Life Technolgies Ltd., Paisley, UK).

### (d) Library preparation, sequencing, processing of NGS data, and variant calling

We prepared Illumina TruSeq PCR free (FC-121-3003; Illumina United, Fulbourn, UK) sequencing libraries with 350 bp insert lengths according to manufacturer’s instructions with slight modifications. We processed the sequencing data files using custom python3 or bash scripts that allowed batch processing on high performance cluster computers. Workflows were based on GATK Best Practices, GATK version 3.6 or higher [25]. The NGS data processing pipeline involved initial processing of raw sequence data, mapping, re-aligning of sequence data around indels, and variant discovery (Supplementary Methods, https://github.com/syllwlwz/Divergence-Scans/tree/master/SNP_calling_arenosa). Neutral population structure was assessed based on initial NGS data processing as conducted for environmental association analysis (EAA), employing the putatively neutral four-fold degenerate sites from the filtered vcfs. We extracted the allele frequency per individual and used this data for a PCA using the R-package FactoMineR [26].

### (e) Genome Scans, large-effect variant identification, candidate gene lists and test for convergent evolution

For each population pair, the genome was partitioned into windows of 25 consecutive SNPs, for which we calculated the pairwise metrics Diversity-Divergence residuals (DD) [27, 28], Wright’s Fixation Index (F_ST_) [28-30], Two-Dimensional Site Frequency Spectrum Composite Likelihood Ratio test (Nielsen 2dSFS) [27, 28, 31], Maximum Absolute Net Divergence (d_xY_) [28, 32], and T(F-LK) *P*-value (Flk) [33], VarLD [34] and the single population metrics Tajima’s *D* [35] and Fay [35] and Wu’s *H* [36], as well as SweeD [31, 37]. Candidate windows for selection were identified as ≥ 99.9%ile for either one of the pairwise metrics with the exception of DD (≤ 0.1%ile) for Klet and Kowa of *A. halleri*. Orthologous *Arabidopsis thaliana* gene identifiers were retrieved for *A. lyrata* genes and genes not assigned to an OrthoGroup were submitted for a local blastx on the nr database. All variants (SNPs and indels) were annotated and their effects predicted by SNPEFF [38] based on the *Arabidopsis lyrata* annotation version 2 [39]. All candidate genes underwent a custom filtering process. The genome-wide output of SnpEff was used to identify large-effect variants at divergent frequencies between populations of a pair to determine additional candidate genes. Additionally, coverage was calculated per gene or per exonic gene content and the number of paralogous genes within the same orthoGroup was extracted. To identify common candidate genes, Venn diagrams of candidate genes were generated with Venny 2.1.0 [40] and hypergeometric tests were performed to compare observed and expected overlaps. A gene function enrichment test was performed for each population pair using the ClueGO app version 2.5.2 [41] in Cytoscape version 3.6.1 [42] using the *A. thaliana* gene identifiers and the gene ontology (GO) “BiologicalProcess”.

### (f) Identifying divergence signatures, Environmental Association Analysis, and compilation of candidate gene lists

To identify divergence signatures, we calculated allele frequency difference (AFD), d_xY_ [28, 32], F_ST_ [28-30], DD [27, 28], and Tajima’s *D* [35] in SNP based windows. We defined a divergence signature as ≥ 99.5%ile windows in the empirical distributions for each metric (https://github.com/SailerChristian/Divergence_Scan). To identify DivergenceScan candidate genes that contain SNPs strictly associated with soil trace metal element (TME) concentration, we tested the divergence scan candidates for association to environmental variables using Bayenv2 [43], which allows for testing in a two-step process. In order to be able to infer effects on protein structure, we selected environmentally very strongly associated SNPs (EA SNPs, BF ≥ 100 [44]) that cause a non-synonymous change according to the annotation using SnpEff [38]. To draw conclusions about convergence, we created a union list of extractable Cd and extractable Zn EA SNPs per contrast and species and identified the intersect between the two contrasts within each species.

### (g) Homology modeling and re-assessment of putative intron-exon boundaries

Homology models of AhCLP1 were generated with Modeller 9.20 [45] using the structures of *Actinidia chinensis* (PDB: 2ACT), *Tabernaemontana divaricata* (PDB: 1IWD), *Zingiber officinale* (PDB: 1CQD) and *Homo sapiens* (PDB: 1BY8, identified using PSIPRED [46] and incorporated to model the pro-peptide). The final model was determined based on DOPE score. Multiple sequence alignments were generated using Clustal Omega (https://www.ebi.ac.uk/Tools/msa/clustalo/) and Modeller 9.20 [45], with colour scheme showing percentage identity through Jalview [47].

Intron-exon boundaries were initially determined by alignment to the annotated *A. lyrata* reference genome. Boundaries were re-defined using the *A. halleri* reference genome [48]. *Arabidopsis halleri* introns from gene Araha.2668s0004 were aligned to the genomic consensus sequences of *AhCLP1.* In this way, intron 2 was expanded by 6 nucleotides at the 3’ end (2 amino acids) and intron 4 was expanded at the 5’ end by 117 nucleotides for the M site allele and 185 nucleotides for the NM site allele.

## 3. Results

### (a) Choice of site pairs and edaphic characterization of sites and microhabitats

While sampling *A. halleri* at 165 European sites [6], we noticed the additional presence of *A. arenosa* plants at a small subset of M and NM *A. halleri* sites. According to soil multielement analysis, *A. halleri* grows in highly metal-contaminated soil patches at M sites whereas *A. arenosa* typically occupies low-metal soil microhabitats (data not shown). However, at Miasteczko Śląskie/PL (Mias) and Kletno/PL (Klet), individuals of both species grew in M soil microhabitats of highly similar composition (figure 1, table S1, S2, dataset S1). Between the two species, the only significant differences were higher extractable Al and extractable Cu concentrations in *A.* halleri-adjacent soil at Klet (A. *halleri* 513 ± 99 mg Al kg^-1^ soil, *A. arenosa* 384 ± 131 mg Al kg^-1^ soil, F_1,16_ = 4.930, *p* < 0.05; *A. halleri* 53.25 ± 30.84 mg Cu kg^-1^ soil, *A. arenosa* 27.72 ± 14.53 mg Cu kg^-1^ soil, F_1,16_ = 5.514, *p* < 0.04; table S2).

**Figure 1.**
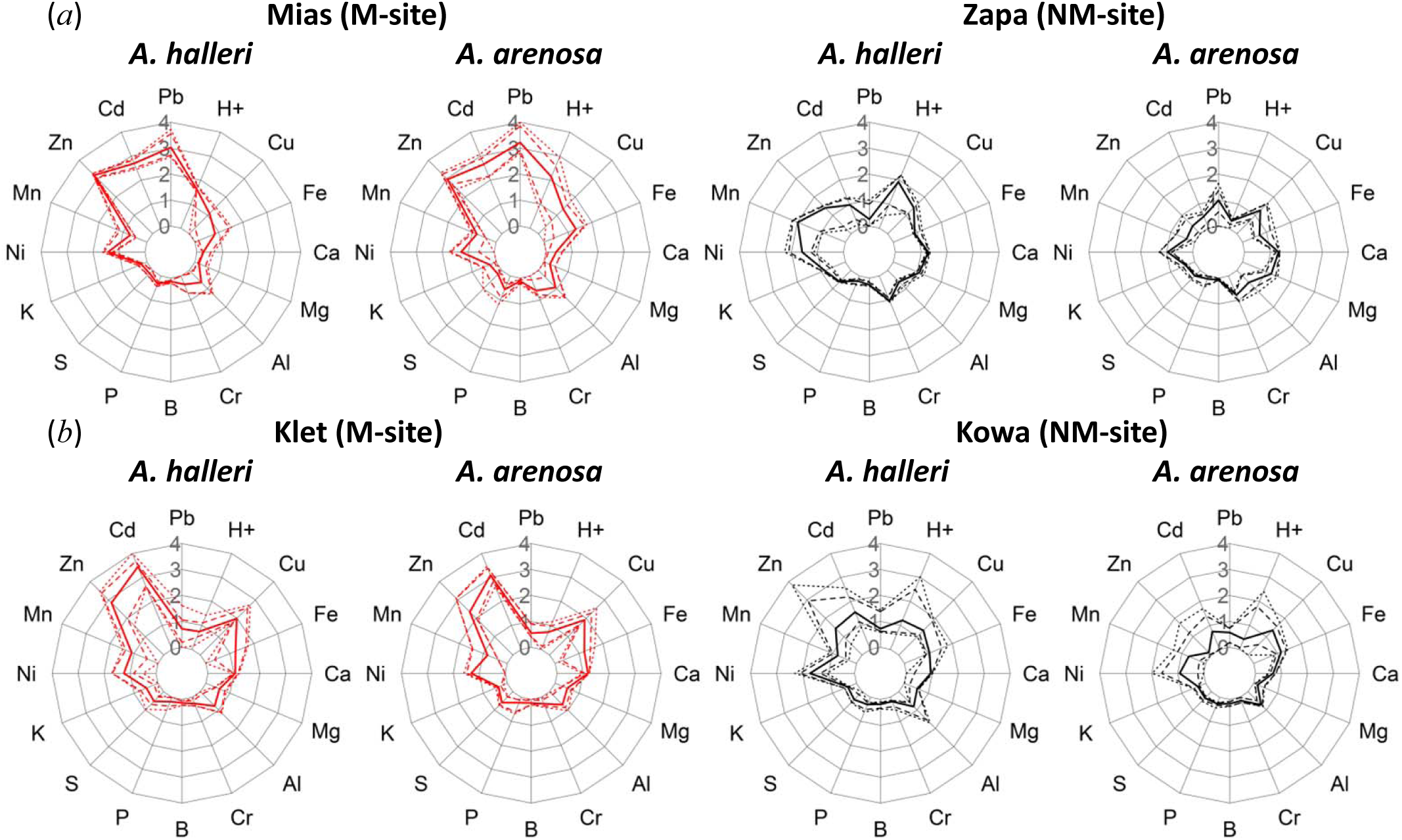
Mineral composition of exchangeable fraction of soils, and soil pH. (*a*) Site pair Miasteczko Śląskie (Mias) and Zakopane (Zapa), (*b*) Site pair Kletno (Klet) and Kowary (Kowa). Concentrations of elements were determined in 0.01 M BaCl_2_ extracts of soils collected in the field directly adjacent to roots of the plant individuals that we resequenced (see Methods). Concentrations [mg element kg^-1^ dry soil mass] were normalized to the global minimum per site pair across both species, and subsequently Log_10_-transformed. Shown are the median (solid line), 10 and 90%iles (dashed lines), minimum and maximum (dotted lines) for each site per species (*n* = 5 to 9 plant individuals) for metalliferous (M, red) and non-metalliferous (NM, black) soils.

Based on these two focal M sites that were both very high in soil exchangeable Zn and Cd, we chose geographically proximal NM sites that also hosted both species, namely Zakopane/PL (Zapa) and Kowary/PL (Kowa). Plant-proximal soils at these NM sites contained tenfold less or lower average exchangeable soil Zn and Cd concentrations (figure 1). Indeed, exchangeable soil cadmium and zinc concentrations differentiated M from NM sites for both species, based on one-way ANOVA (linear model; Cd: F_3,26_ = 149.7, *p* = 2.2*10^-16^; Zn: F_3,26_ = 68.63, *p* = 1.8*10^-12^; table S3). These major global contrasts between M and NM soils were also evident in the extractable fraction of soils (figure S1). Thus, we were able to address the parallel evolution of edaphic adaptation to calamine M soil through the comparison between the site pairs Mias-Zapa and Klet-Kowa in both species, as well as through the comparison between species for either of the two site pairs. Between species, soil mineral composition differed more at the NM sites, with generally lower soil pH and higher soil Zn, Cd and Ni in *A. halleri* microhabitats compared to those of *A. arenosa.*

### (b) Experimental test for adaptation to metalliferous soil

Under climate-controlled growth chamber conditions survival of *A. halleri* was 100% irrespective of plant origin and experimental soil treatment (figure 2a; figure S2, table S4). Similarly, there was 100% survival of *A. arenosa* plants on control soil irrespective of plant origin. However, on M Mias soil, only *A. arenosa* of Mias origin were able to survive (100% survival rate), whereas all plants originating from Zapa died (survival rate 0%) (figure 2*b*). Biomass production of *A. halleri* exhibited crossing reaction norms, which indicates local adaptation of Mias plants to Mias soil (figure 2*c*, figure S2, significant interaction between plant genotype and treatment soil at *p* = 0.013, table S4). Similarly *A. arenosa* exhibited a signature of local adaptation (figure 2d, *p* < 0.001). The observed trends were reproduced in an independent experiment (figure S3, table S4).

**Figure 2.**
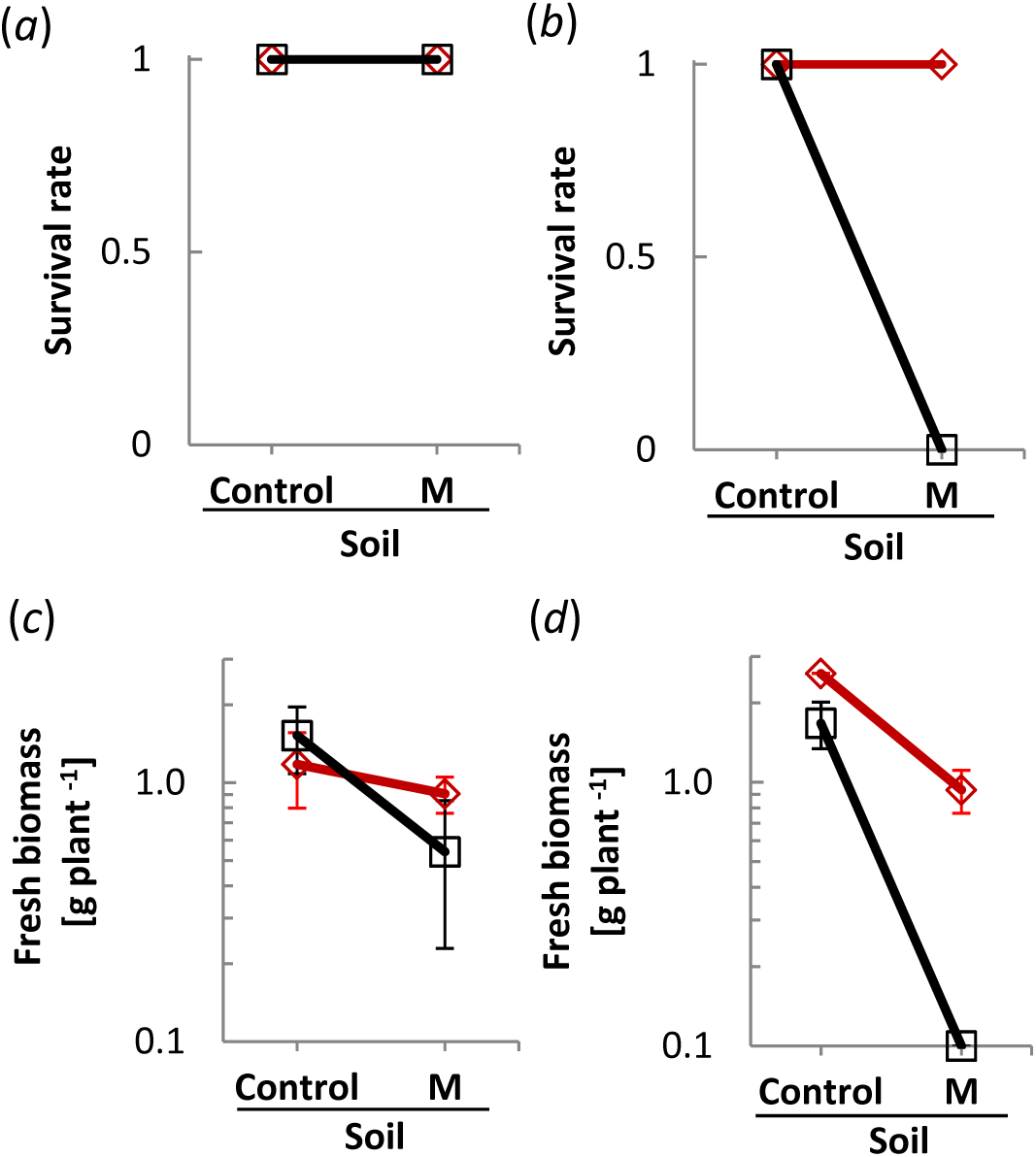
Experimental test for adaptation to metalliferous soil. (*a*,*b*) Survival of *A. halleri* (*a*) and *A. arenosa* (*b*) plants originating from Mias (M site, red) and Zapa (NM site, black) transferred into metalliferous (Mias) or non-metalliferous (control) soil. (c,d) Fresh biomass of *A. halleri* (*c*) and *A. arenosa* (*d*) plants originating from Mias (M site, red) and Zapa (NM site, black) transferred into metalliferous (Mias) or non-metalliferous (control) soil. Shown are means and standard deviations of survival rate (*a*,*b*) and fresh above-ground biomass (*c*,*d*) after 6 weeks of cultivation on experimental soils (see table S4 for details).

### (c) All population pairs are genetically distinct

To assess population structure we conducted a principal component analysis (PCA) using putatively neutral (four-fold degenerate) sites. We found that the *A. halleri* individuals from Mias and Zapa were genetically more closely related to one another than those at Klet and Kowa, whereas in *A. arenosa* we observed the opposite (figure S4*a*,*b*). Furthermore, for *A. halleri* the first principal component separated individuals from Klet from those at the other three populations (figure S4*a*). This is most likely caused by a relative excess of low frequency variants in Klet, as illustrated by folded site frequency spectra (figure S4*e*), and consistent with a population bottleneck at Klet. Importantly, the neutral population structure was clearly distinct from the dominant contrast between M and NM soil types for both species.

### (d) Population pair-wise identification of candidate genes for selection at metalliferous sites

To obtain candidate gene-coding loci underlying repeated adaptation to M soils, we next scanned genomes sampled from these populations for selective sweep signatures. We determined gene content of candidate 25-SNP windows based on unique *A. lyrata* gene identifiers, and filtered candidate gene-coding loci according to stringent criteria (see Methods). In *A. halleri*, this identified 94 candidate gene-coding loci based on any one metric in Mias relative to Zapa (figure 3, dataset S2). Independently, we identified 81 genes exhibiting predicted large-effect SNPs and 74 genes exhibiting predicted large-effect indels (see Methods; dataset S2). In Klet by comparison to Kowa, we identified 73 candidates in genome scans (figure 3), as well as 488 large-effect SNP-and 379 large-effect indel-containing genes (dataset S3). In *A. arenosa* at Mias compared to Zapa, we identified 135 candidate genes in divergence window-based scans, and 15 and 33 genes containing predicted high-effect SNPs and indels, respectively (figure 3, dataset S4). Finally, in *A. arenosa* at Klet compared to Kowa, we identified 147 candidate genes, as well as 5 and 16 genes containing high-effect SNPs and indels, respectively (figure 3, dataset S5). We conducted an enrichment analysis on GO “biological pathways” with Cytoscape for the candidate genes identified (see Methods; datasets S2 – S5). In *A. halleri*, this identified an overrepresentation among candidates in the functions “Ammonium ion metabolic process” for Mias (*vs.* Zapa) and “Acceptance of pollen” for Klet (*vs*. Kowa), among others (figure 3*b*). In *A. arenosa*, “regulation of sequestering of zinc ion” were most over-represented among candidate genes at Mias (*vs*. Zapa), and “Vacuolar sequestering” for Klet (*vs*. Kowa) (figure 3*b*).

**Figure 3.**
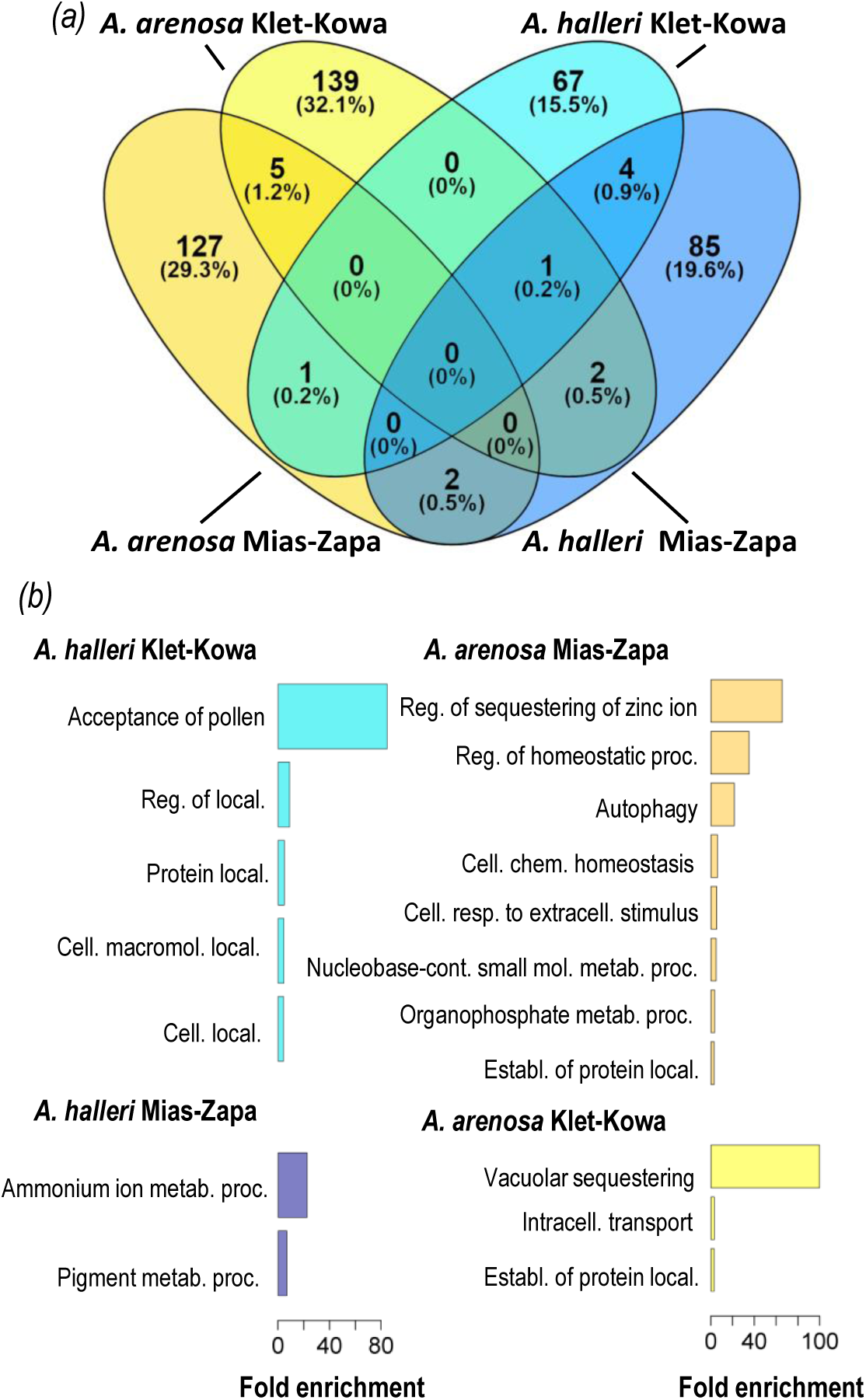
Candidate genes exhibiting signatures of selective sweeps, and functional enrichment analysis. (*a*) Venn diagram shows the number of genes for each population pair in *A. halleri* and in *A. arenosa*, and their intersecting sets. Candidate genes were among the ≥ 99.9%iles for any one pairwise genome scan metric and subsequently filtered manually as described in Methods. (*b*) Gene ontology (GO) biological process annotations enriched candidate genes for each population pair. Shown are GO biological processes (level 1-3) with at least 3-fold overrepresentation (*p* < 0.05) among the same candidates as in (*a*) in comparison to the genome-wide average based on *A. thaliana* orthologues for Mias-Zapa in *A. halleri* (dark blue), in *A. arenosa* (dark yellow), Klet-Kowa in *A. halleri* (light blue) and in *A. arenosa* (light yellow).

### (e) Degree of convergent evolution

To identify candidate genes undergoing convergent selection, we identified intersecting sets of genes from each genome scan between the two site pairs and between the two species. In a conservative approach, we identified 5 candidate genes exhibiting selective sweep signatures that were convergent between both population pairs Mias (*vs*. Zapa) and Klet (*vs*. Kowa) in *A. halleri* and also *A. arenosa* (both *p* < 0.001, hypergeometric test; figure 3*a*, table 1). There was no convergent candidate gene across both site pairs common to both species (see dataset S6 for a less conservative gene list that additionally includes those candidate genes identified through the presence of large-effect SNPs or indels at high allele frequencies). One of the convergent candidate genes across both site pairs in *A. halleri* (AL8G20240, annotated as tRNA dihydrouridine synthase, see table 1, figure S5*c*) was also a candidate for selection in Klet (*vs*. Kowa) in *A. arenosa*. Two candidate genes were in common between the two species at Mias (*vs*. Zapa), and one gene at Klet (*vs*. Kowa) (n.s., hypergeometric test). Three genes were candidates at Mias in *A. halleri* and at Klet in *A. arenosa* (*p* < 0.01), and one gene was a candidate at Mias in *A. arenosa* and at Klet in *A. halleri* (n.s.). Additionally, there was some convergence among gene functional categories related to cellular protein localization (figure 3*b*). These were significantly overrepresented at Klet (**vs*.* Kowa) in *A. halleri* and at both site pairs in *A. arenosa* (protein localization, establishment of protein localization, regulation of protein localization, cellular macromolecule localization, cellular localization, intracellular transport; figure 3*b*).

**Table 1.** Convergent candidate genes for selection as identified in this study.

| *A. lyrata*<br>Genome<br>Identifier | *A. thaliana*<br>Genome<br>Identifier | Short<br>Gene<br>Name | Gene Annotation |
| --- | --- | --- | --- |
| ***Arabidopsis halleri* candidate genes convergent between site pairs** |  |  |  |
| AL2G22120 | AT1G65430 | ARI8 | Ariadne 8; ubiquitin protein ligase<sup>s</sup> |
| AL2G31400 | AT1G71820 | SEC6 | Exocyst complex gene family member; vesicle secretion |
| AL5G40920 | AT3G59040 |  | Tetratricopeptide repeat (TPR)-like superfamily protein |
| AL5G40930 | AT2G43020 | PAO2 | Polyamine oxidase 2<sup>s</sup> |
| AL8G20240 | AT5G47970 |  | tRNA dihydrouridine synthase; Aldolase-type TIM barrel family* |
| ***Arabidopsis arenosa* candidate genes convergent between site pairs** |  |  |  |
| AL2G12590 | AT1G63010 | PHT5;1 | Major Facilitator Superfamily, SPX domain; vacuolar Pi sequestration<sup>s</sup> |
| AL2G30820 | AT1G71210 |  | Pentatricopeptide repeat (PPR) superfamily protein<sup>g,r</sup> |
| AL7G17030 | AT4G34260 | AXY8 | Altered xyloglucan 8; 1,2-α-L-fucosidase; mutant is Al-tolerant<sup>g,r</sup> |
| AL8G14060 | AT5G45140 | NRPC2 | nuclear RNA polymerase C2<sup>g,f</sup> |
| AL7G35050 | AT4G19050 |  | NB-ARC protein; GWAS association with H<sub>2</sub>O<sub>2</sub> tolerance |
| **Candidate genes at Mias (vs. Zapa) convergent between species** |  |  |  |
| AL5G26930 | AT3G47640 | PYE | Popeye; bHLH transcription factor acting in iron homeostasis<sup>*,g</sup> |
| AL5G26940 | AT3G47650 | BDS2 | Bundle sheath defective 2; DnaJ/Hsp40 cys-rich domain protein<sup>*,l</sup> |
| **Candidate genes at Klet (vs. Kowa) convergent between species** |  |  |  |
| AL8G20240 | AT5G47970 |  | tRNA dihydrouridine synthase; Aldolase-type TIM barrel family* |
| **Candidate genes convergent between *A. halleri* at Mias (vs. Zapa) and *A. arenosa* at Klet (vs. Kowa)** |  |  |  |
| AL1G26370 | AT1G14470 |  | Pentatricopeptide repeat (PPR) superfamily protein<sup>f,g</sup> |
| AL8G20240 | AT5G47970 |  | tRNA dihydrouridine synthase; Aldolase-type TIM barrel family* |
| AL6G35310 | AT5G23980 | FRO4 | ferric reduction oxidase 4; root surface Cu(II) chelate reductase<sup>r,l</sup> |
| **Candidate genes convergent between *A. halleri* at Klet (vs. Kowa) and *A. arenosa* at Mias (vs. Zapa)** |  |  |  |
| AL1G53790 | AT1G47560 | SEC3B | Exocyst complex gene family member; vesicle secretion |
| ***A. halleri* candidate genes convergent between site pairs identified by EAA** |  |  |  |
| AL1G34900 | AT1G21722 |  | unknown transmembrane protein<sup>f</sup> |
| AL3G38930 | AT3G24250 |  | glycine-rich protein<sup>s</sup> |
| AL3G53910 | AT2G20800 | NDB4 | NAD(P)H dehydrogenase B4<sup>p</sup> |
| AL5G24630 | none |  | Blast: Alpha-D-xyloside xylohydrolase |
| AL6G23510 | none |  | DNA damage repair/tolerance protein DRT100-related (AT5G12940)<sup>r</sup> |
| AL8G32870 | none | CPL1 | Cysteine Protease-like 1 |
| AL8G32880 | none |  | hypothetical protein AXX17-related (AT5G56200)<sup>s</sup> |

\*Large-effect SNPs or indels of allele frequency difference > 0.9 in at least one population pair.
Tissue-specific or strongly enhanced gene expression: <sup>g</sup>germination, <sup>s</sup>siliques, <sup>r</sup>root, <sup>f</sup>flower, <sup>p</sup>pollen

### (f) Environmental association analysis (EAA)

As a complementary approach, we conducted an environmental association analysis (EAA) across all four sites separately in each species. This analysis was justified by our analysis of neutral population structure and by the differentiation in soil composition (see figure 1, figure S4). We thus identified candidate gene-coding loci positioned in a 25-SNP-window exhibiting a signature of relative divergence and containing at least one derived SNP highly associated with the differentiating soil concentrations of exchangeable Cd and/or Zn (DivergenceScan, https://github.com/SailerChristian/Divergence_Scan; see Methods and Supplementary methods). Out of approximately 100,000 and 540,000 SNP positions tested for *A. halleri* and *A. arenosa* (see Table S1), respectively, between 0.047% and 1.12% were identified to exhibit a very strong environmental association (BF ≥ 100). Among these, we additionally required that a candidate gene-coding locus must contain at least one associated non-synonymous SNP and that the environmentally associated SNP must be present at a higher frequency in both metallicolous populations (Klet and Mias) compared to both non-metallicolous populations. In *A. halleri*, a total of seven loci fulfilled all these stringent criteria, whereas for *A. arenosa*, no gene-coding locus was retained (table 1). None of the seven identified genes is associated with a pathway based on the NCBI biosystems repository. Thus, based on this stringent EAA analysis, we did not find a single locus to be convergent between both species.

### (g) Integrating results from both approaches

One candidate gene, *Cysteine Protease-like 1* (AL8G32870, *CPL1*), was identified in *A. halleri* by both i) intersecting results of genome scans combined with potentially selected large-effect SNPs/indels, and ii) EAA of candidate genes in both M populations (table 1, datasets S2 and S3, figure 4). The signature of selection over the candidate gene AL8G32870 was stronger for Mias-Zapa (≥ 99.9%ile for F_st_, d_xY_, Flk, highly negative Tajima’s *D*, ≤ 0.1% for DD) than for Klet-Kowa (figure 4*a*). *AhCPL1* was identified as a candidate gene in the Klet-Kowa population pair based on a large-effect indel exhibiting at high allele frequency difference (allele frequency difference of 0.86, dataset S3). For the initial step in EAA (selecting the candidtes to be tested), F_st_ and d_xY_ were in the ≥ 99.5%ile and DD in the ≤ 0.5%ile. SweeD indicated a strong signal for the Klet, but not the Kowa population. Extreme values of metrics pinpointed exons one to four (figure 4*a*), in agreement with EAA which identified 50% of the significantly associated SNPs in this region (figure 4*b*, *c*). A closer inspection of haplotypes revealed that the predicted proteins at Mias and Klet share amino acid variants at 12 out of a total of 16 variable positions of the predicted protein that differentiate them from Zapa and Kowa (figure S6). Of these 12 predicted amino acids characteristic of M sites, 9 are derived compared to the *A. lyrata* reference.

**Figure 4.**
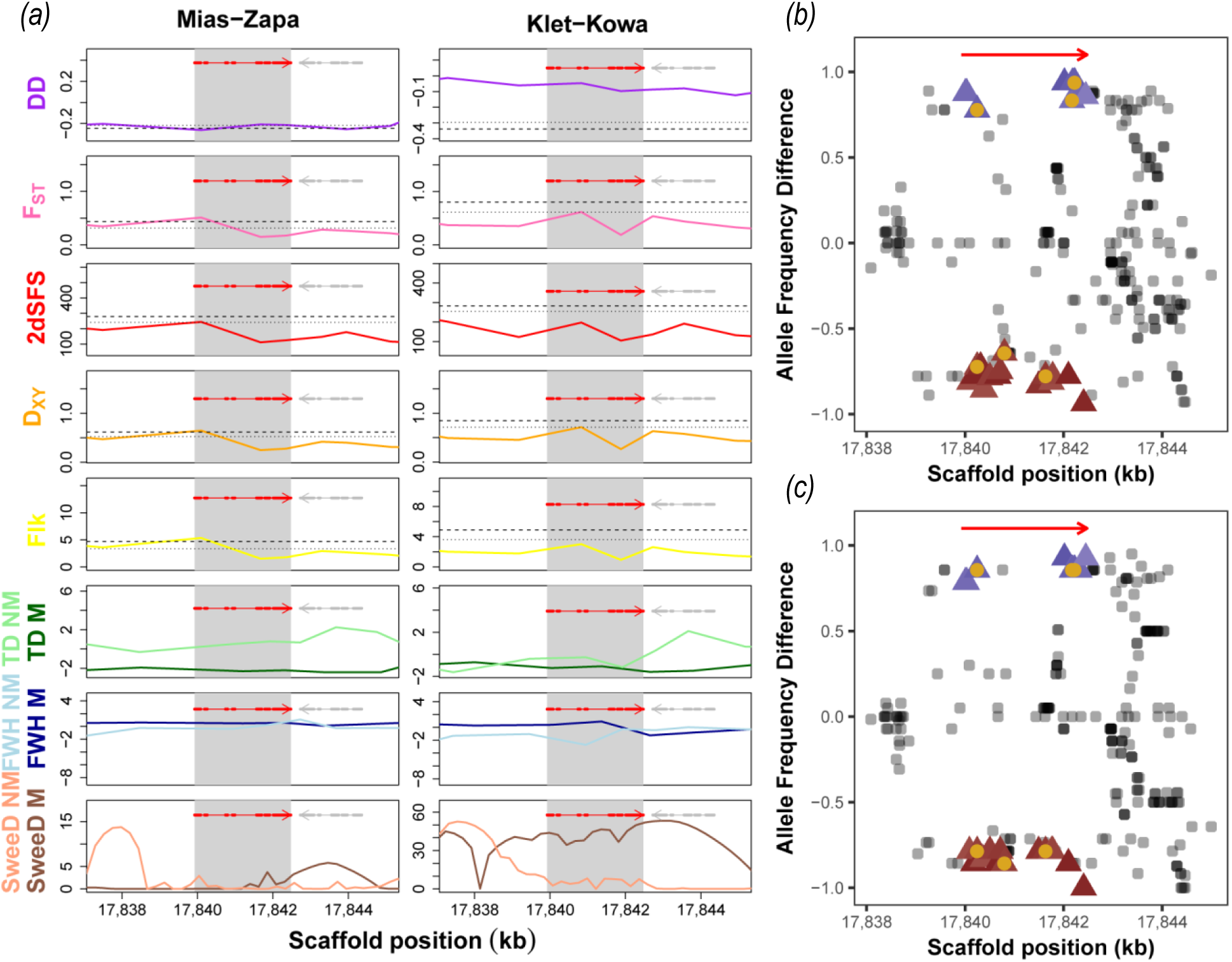
Evidence for convergent selection at the *A. halleri* candidate locus *Cysteine Protease-like 1 (CPL1,* AL8G32870). (*a*) Genome scans. Lines connect datapoints, each representing the centre of a 25-SNP window, for diagnostic metrics for Mias **vs*.* Zapa (left) and Klet *vs*. Kowa (right). The dashed/dotted lines mark the 99.9%/99.5% genome-wide percentiles (0.1%/0.5% for DD), respectively. Red thick/thin lines reflect exons/introns of candidates. (*b*, *c*) EAA results for *CPL1* (red arrow) for Mias-Zako (*b*) and Klet-Kowa (*c*). Each datapoint marks the position of one SNP and its allele frequency difference between the metallicolous and non-metallicolous population. Blue/red triangles represent SNPs positively/negatively correlated and very strongly associated (BF ≥ 100) with exchangeable Cd concentrations in soil, with non-synonymous variants marked by yellow circles (identical results for exchangeable zinc concentrations; not shown). More negative DD residuals (in a) indicate lowered diversity relative to the degree of between-population differentiation; the magnitude of F_st_ values reflects the degree of between-population relative differentiation; raised 2dSFS values reflect a shift in the two-dimensional site frequency spectrum consistent with positive selection; d_xY_ quantifies absolute net between-population absolute divergence; Flk indicates the degree of population differentiation adjusted for relatedness. A negative Tajima’s *D* (TD) reflects an excess of low frequency variants; a negative Fay and Wu’s H (FWH) an excess of high-frequency derived SNPs; elevated SweeD represents a shift in the site frequency spectrum indicating a selective sweep (see Methods). Metallicolous (M: Mias, Klet)/ non-metallicolous (NM) populations are shown in dark/light colour in (*a*).

### (h) Homology modeling of AhCPLI proteins

To gain insight into possible consequences of the amino acid exchanges in the predicted *Ah*CPL1 Papain-like cysteine protease protein, we generated a homology model of the protein structure. An initial multiple sequence alignment (figure S7*a*) suggested that a segment of the protein was missing (t2, *A. lyrata* genome v2.1, JGI Phytozome, https://phytozome.jgi.doe.gov/pz/portal.html). Intron-exon boundaries were then reassessed based on *A. halleri* v1.1 (JGI Phytozome). We hypothetically expanded the 5’- end of the fifth exon by 177 nucleotides (59 amino acids), leading to the incorporation of the missing protein motifs in the metallicolous *Ah*CPL1 variants (figures S7*b*, S8). Through this alteration in intron-exon boundaries a frameshift arising through an 8-bp insertion renders the predicted non-metallicolous *Ah*CPL1 variants non-functional. In the structures generated by homology modelling the protein consists of an N-terminal 42-amino acid propeptide, which is likely to be cleaved during enzyme maturation, and the mature protein consisting of characteristic L- and R-domains characterized by alpha helices and beta sheets, respectively (figure S9, N-terminus included). While the protein is relatively divergent from other proteins with available cysteine protease structures, the active site is recognizable in the model and contains all four catalytically active residues in close proximity (Q62, C68, H212 and N234).

## 4. Discussion

The objective of this study was to probe for candidate genes that have undergone convergent selection on calamine M soils. We focused on genes with evidence for selection at two M sites, each of them in comparison to a geographically proximal NM site, in the two closely related and genetically tractable species *A. halleri* and *A. arenosa* (table S1). This sampling design was chosen in order to gain greater power and test for convergence between species. We analysed genome-wide resequencing data obtained from field-collected individuals in relation to the mineral composition of root-proximal soils from the same individuals. Our objective of identifying convergent genes thus targeted selection by environmental factors common among (rather than specific to) population pairs.

The two M sites Mias and Klet were previously known for their vegetation type characteristic of calamine M soils [6]. In agreement with this, we found highly elevated Zn and Cd levels in root-proximal soils which distinguish both of these sites from the our NM sites (figure 1, tables S2 and S3, dataset S1). Our hypothesis of local adaptation to Mias soil in both species was supported by differing plant biomass production depending on population and soil type in independent experiments (figure 2, table S4; figures S2 and S3). In particular, upon cultivation on Mias soil, plants of NM origin were more severely affected in *A. arenosa* than in *A. halleri*. Indeed, it is well-known that *A. halleri* exhibits enhanced tolerance to Zn and Cd species-wide [12, 49], which is exceedingly rare [5, 6]. Thus, local adaptation to M soil at Mias, and likely also to Klet, must confer a larger degree of additional heavy metal tolerance to *A. arenosa* than to *A. halleri.*

Genome scans identified few candidate genes being convergently under selection at both site pairs in either of the two species (figure 3). Similarly, the number of genes convergent between *A. arenosa* at Klet (*vs*. Kowa) and *A. halleri* at Mias (*vs*. Zapa) was very low, but both also exceeded the number expected by chance alone (Table S5). This latter finding was consistent with our expectations based on species-wide traits as outlined above, and with the fact that Zn and Cd concentrations were considerably lower in Klet soils than in Mias soils (dataset S1).

At the level of gene ontologies, protein localization was common to three out our four contrasts and thus a candidate function under convergent selection (figure 3*b*). To date, functions in cellular protein trafficking are neither known as vulnerable targets of Zn or Cd toxicity nor as a means of attaining cellular metal tolerance. Especially the cellular targets of heavy metal toxicity are poorly understood to date.

Among the candidate genes exhibiting some degree of convergence, we identified genes known to act in the homeostasis of Fe (bHLH transcription factor-encoding *PYE* [50]) and Cu (cell surface Cu(II) chelate reductase-encoding *FRO4* [51]). Homeostasis of essential nutrients Fe and Cu is a well-known target of Zn and Cd toxicity [52-56] (table 1, figure S5). These data suggest a modest degree of biologically relevant convergent evolution, more prominently within species and between sites, but also between species across environmental contrasts of comparable magnitude. Predicted functions of other convergent candidate genes were diverse, generally poorly characterized, and sometimes in a context of known relevance under heavy metal stress, for example cell wall composition or DNA damage repair [57, 58].

Based on existing knowledge of the molecular basis of species-wide metal hypertolerance, we had expected to identify candidate genes with direct roles in the detoxification of Cd^2+^ and Zn^2+^, such as their transmembrane transport and binding [11, 21,59]. This was indeed the case in *A. arenosa* at one site pair (Mias-Zapa), yet with no apparent convergence (figure 3*b*). The genes underlying within-species variation in metal tolerance of plants are unknown. Almost all genes that have been experimentally demonstrated to contribute to naturally selected species-wide Zn and Cd hypertolerance, and also many candidate genes implicated in these traits, are copy number-expanded in *A. halleri*, with paralogues almost identical in sequence [59-61]. Genes with such sequence properties are usually disregarded in genome scans by excluding both reads mapping to multiple loci in the genome and genomic regions with excessive short read coverage. Beyond these technical issues, it can be very difficult to detect selective sweeps at such loci, because they commonly exhibit complex patterns of polymorphism resulting from ectopic gene conversion or illegitimate recombination. This was exemplified by the copy number-expanded metal hyperaccumulation and metal hypertolerance locus *HMA4* of *A. halleri* [62].

Through both genome scan and EAA approaches, we identified the *A. halleri* locus corresponding to AL8G32870 in the *A. lyrata* reference genome as a convergent candidate gene between both population pairs (table 1, figure 4, datasets S6 and S7). In confirmation, we observed convergent high frequency non-synonymous sequence divergence at this locus specific to M sites (figure S6). According to between-population differentiation metrics, this gene was thus not among the strongest selective sweep candidates in Klet-Kowa, but instead entered the list of candidates through an indel observed with a high between-population allele frequency difference (dataset S3). *AhCPL1* is predicted to encode a papain family cysteine protease that has no orthologue in *A. thaliana* (figure S7). An adjustment of the intron-exon boundaries raised the possibility that a functional CPL1 protein may be specific to the M sites Mias and Klet (figure S8). *AhCPL1* is the first candidate metal hypertolerance gene of a Brassicaceae lacking a homologue in a syntenic position in the *A. thaliana* genome. This finding may contribute to explaining why no single *A. thaliana* accession was identified to grow naturally on a M soil.

The possible molecular role of AhCPL1 in metal tolerance remains unknown. The papain family cysteine protease genes of *A. thaliana Response to Dehydration 19* and *21 (RD19, RD21)* have long been known as components of the transcriptional response to dehydration and salt stress [63]. More recently, RD19 was reported to function in signalling to trigger anti-microbial defences [64]. In wheat, a cysteine protease was transcriptionally upregulated under aluminium stress [65]. In *Chlamydomonas* sp., oxidative stress induced a cysteine protease which provided Cd tolerance [66]. In *A. halleri*, transcript levels encoding a putative cysteine proteinase (AT2G27420) were observed to be about 30-fold higher than in *A. thaliana* according to microarray-based cross-species transcriptomics [67].

It must be kept in mind that there were several important environmental factors differing between sites. Specifically, when compared to the NM soil, Mias soil was between 3- and 10-fold lower in exchangeable Ca^2+^, a nutritional condition that is well known to enhance the toxicity of divalent heavy metal cations [68]. Mias soil was also lower in several nutrients, i.e. exchangeable K+, Mg^2^+ and Mn^2^+, in contrast to the Klet-Kowa pair (dataset S1, table S2). Importantly, Mias soils were on average more than 100-fold higher in exchangeable Pb, a heavy metal with extremely high toxicity potential, than Zapa soils, whereas soil exchangeable Pb was not elevated at Klet (*vs*. Kowa). Conversely, Klet soils contained about 4-fold elevated exchangeable concentrations of Cu, which can be highly toxic to plants. As a former uranium mine, Klet soil may additionally contain elevated levels of toxic decay products of uranium that were not quantified here (e.g. polonium, thallium). Consequently, the limited number of convergent candidate genes identified here is unsurprising in relation to the multi-factorial stress of local soil environments (figure 3, datasets S2 to S7).

The candidates obtained through the sequence divergence-based approaches pursued in this study, and their intersection, overall suggested a limited sensitivity of these approaches. Nevertheless, we could identify several candidate genes convergent between site pairs and between species, as well as convergent sequence variants in one convergent candidate gene. Species-specific biology and metal handling may explain the small number of convergent candidates identified between species. However, our data suggest the existence of functional gene network convergence, but with differing proximal molecular factors mediating functional convergence. Considerable future effort will be required for the functional characterization of identified candidate genes and networks. Additionally, a greater degree of convergent evolution within species may be observed in future work at M sites chosen for higher similarity in soil composition, whereas here we chose M sites exclusively based on the presence of both species.

## List of Supplementary Files

### Supplementary Methods

**Figure S1.** Mineral composition of extractable fraction of soils, and soil pH.

**Figure S2.** Photographs of plants in experiment testing for plant adaptation to metalliferous soil.

**Figure S3.** Independent experiment addressing local adaptation to metalliferous soil.

**Figure S4.** Population structure and site frequency spectra.

**Figure S5.** Evidence for selection on convergent genes.

**Figure S6.** Alignment of the predicted variants of the *Arabidopsis halleri* Cysteine Protease-Like 1 (CPL1) protein.

**Figure S7.** Multiple sequence alignment (MSA) of Papaine-like cysteine proteases.

**Figure S8.** Multiple sequence alignment (MSA) of consensus genomic DNA sequence at the *AhCPL1* locus.

**Figure S9.** Predicted structure of the Mias AhCPL1 protein variant shown as a ribbon.

**Table S1.** Number of *Arabidopsis arenosa* and *Arabidopsis halleri* individuals used for the genome scans

**Table S2.** Statistical analysis of soil data

**Table S3.** Statistical analysis of soil data for EAA.

**Table S4.** Experiments testing for plant adaption to metalliferous soil.

**Table S5.** Hypergeometric tests for the number of overlapping genes expected by chance.

**Dataset S1.** Soil pH and concentrations of mineral elements in extractable and exchangeable fractions of soil adjacent to plant individuals sampled in the field.

**Dataset S2.** Candidate genes Mias *vs*. Zapa in *Arabidopsis halleri*.

**Dataset S3.** Candidate genes Klet *vs*. Kowa in *Arabidopsis halleri*.

**Dataset S4.** Candidate genes Mias *vs*. Zapa in *Aarabidopsis arenosa*.

**Dataset S5.** Candidate genes Klet *vs*. Kowa in *Aarabidopsis arenosa*.

**Dataset S6.** Comprehensive list of convergent genes including filtered candidate genes from genome scans and from searches for large-effect indels and single nucleotide polymorphisms.

**Dataset S7**. Convergent genes in *Arabidopsis halleri*, based on DivergenceScan and environmental association analyses.

### Data accessibility

Sequence data that support the findings of this study have been deposited in the Sequence Read Archive (SRA; https://www.ncbi.nlm.nih.gov/sra) with the primary accession code PRJNAxxxxxx (available upon publication of this manuscript at http://www.ncbi.nlm.nih.gov/bioproject/xxxxxx]

### Authors’ contributions

U.K., C.S, and L.Y. conceived the research. U.K., V.P., C.S., L.S. and L.Y. designed experiments and data analysis. V.P. and C.S. performed experiments. S.B., V.P., C.S. and L.S. analyzed the data. All authors wrote and edited the manuscript.

### Funding

This work was funded by Research Priority Programmes SPP1529 ADAPTOMICS (Kr1967/10-1 and −2 to U.K.) and SPP1819 RAPID EVOLUTION (Kr1967/16-1 to U.K.). L.Y. acknowledges funding from the European Research Council (ERC) under the European Union’s Horizon 2020 research and innovation programme (grant agreement ERC-StG 679056 HOTSPOT) and the UK Biological and Biotechnology Research Council (BBSRC) via grant BB/P013511/1 to the John Innes Centre. U.K. acknowledges funding from the ERC (grant agreement ERC-AdG 788380 LEAP-EXTREME). C.S. was funded by an SNSF Early Postdoc.Mobility fellowship (P2ZHP3_158773).

## Acknowledgements

We thank Petra Düchting for element analysis, Justin Anderson (now Nunhems, The Netherlands) for generating orthoGroups, Andreas Aufermann and Jan Riering for plant cultivation, Aitor Gonzaga Moltó (now DSMZ, Braunschweig, Germany) for plant sampling (all Ruhr University Bochum). We are grateful to the CeBiTec – Center for Biotechnology Bielefeld-Gießen Center for Microbial Bioinformatics (BiGi) of the BMBF-funded German network for bioinformatics infrastructure (de.NBI, grant 031A533) for provision of compute resources and general support. This research was supported in part by the NBI Computing infrastructure for Science (CiS) group.

## References

1. Nieboer, E. & Richardson, D. H. S. 1980 The replacement of the nondescript term “heavy metals” by a biologically and chemically significant classification of metal ions. Environ Poll Series B - Chem Phys 1, 3-26.

2. Brady, K. U., Kruckeberg, A. R. & Bradshaw, H. D. J. 2005 Evolutionary ecology of plant adaptation to serpentine soils. Annu. Rev. Ecol. Evol. Syst. 36.

3. Wójcik, M., Gonnelli, C., Selvi, F., Dresler, S., Rostański, A. & Vangronsveld, J. 2017 Chapter One – Metallophytes of Serpentine and Calamine Soils – Their Unique Ecophysiology and Potential for Phytoremediation. In Advances in Botanical Research (eds. A. Cuypers & J. Vangronsveld), pp. 1-42, Academic Press.

4. Ernst, W. H. O. 1974 Schwermetallvegetationen der Erde. Stuttgart, Germany, Gustav Fischer Verlag.

5. Bert, V., MacNair, M. R., De Laguérie, P., Saumitou-Laprade, P. & Petit, D. 2000 Zinc tolerance and accumualtion in metallicolous and non metallicolous populations of *Arabidopsis halleri* (Brassicaceae). New Phytol 146, 225-233.

6. Stein, R. J., Höreth, S., de Melo, J. R., Syllwasschy, L., Lee, G., Garbin, M. L., Clemens, S. & Krämer, U. 2017 Relationships between soil and leaf mineral composition are element-specific, environment-dependent and geographically structured in the emerging model *Arabidopsis halleri*. New Phytol 213, 1274-1286. (D0I:10.1111/nph.14219).

7. Przedpelska, E. & Wierzbicka, M. 2007 *Arabidopsis arenosa* (Brassicaceae) from a lead-zinc mine waste heap in southern Poland. Plant Soil 299, 43-53.

8. Turisova, I., Strba, T., Aschenbrenner, S. & Andras, P. 2013 *Arabidopsis arenosa* (L.) Law. on metalliferous and non-metalliferous sites in central Slovakia. Bull Environ Contam Toxicol 91, 469-474. (D0I:10.1007/s00128-013-1074-8).

9. Arnold, B. J., Lahner, B., DaCosta, J. M., Weisman, C. M., Hollister, J. D., Salt, D. E., Bomblies, K. & Yant, L. 2016 Borrowed alleles and convergence in serpentine adaptation. Proc Natl Acad Sci U S A 113, 8320-8325. (D0I:10.1073/pnas.1600405113).

10. Turner, T. L., Bourne, E. C., Von Wettberg, E. J., Hu, T. T. & Nuzhdin, S. V. 2010 Population resequencing reveals local adaptation of *Arabidopsis lyrata* to serpentine soils. Nat Genet 42, 260-263.

11. Krämer, U. 2010 Metal hyperaccumulation in plants. Annu Rev Plant Biol 61, 517-534.

12. Meyer, C. L., Juraniec, M., Huguet, S., Chaves-Rodriguez, E., Salis, P., Isaure, M. P., Goormaghtigh, E. & Verbruggen, N. 2015 Intraspecific variability of cadmium tolerance and accumulation, and cadmium-induced cell wall modifications in the metal hyperaccumulator *Arabidopsis halleri*. J Exp Bot 66, 3215-3227. (DOI: 10.1093/jxb/erv144).

13. Meyer, C. L., Kostecka, A. A., Saumitou-Laprade, P., Creach, A., Castric, V., Pauwels, M. & Frerot, H. 2010 Variability of zinc tolerance among and within populations of the pseudometallophyte species *Arabidopsis halleri* and possible role of directional selection. New Phytol 185, 130-142. (DOI:10.1111/j.1469-8137.2009.03062.x).

14. Clemens, S. 2001 Molecular mechanisms of plant metal tolerance and homeostasis. Planta 212, 475-486. (DOI:10.1007/s004250000458).

15. Peer, W. A., Mahmoudian, M., Freeman, J. L., Lahner, B., Richards, E. L., Reeves, R. D., Murphy, A. S. & Salt, D. E. 2006 Assessment of plants from the Brassicaceae family as genetic models for the study of nickel and zinc hyperaccumulation. New Phytol 172, 248-260. (DOI:10.1111/j.1469-8137.2006.01820.x).

16. Szarek-tukaszewska, G. & Niklińska, M. 2002 Concentration of alkaline and heavy metals in *Biscutella laevigata* L. and *Plantago lanceolata* L. growing on calamine spoils (S. Pland). Acta Biologica Cracoviensia Series Botanica 44, 29-38.

17. Szarek-tukaszewska, G. & Grodzińska, K. 2011 Grasslands of a Zn-Pb post-mining area (Olkusz ore-bearing region, S Poland). Polish Botanical Journal 56, 245-260.

18. Kenderesova, L., Stanova, A., Pavlovkin, J., Durisova, E., Nadubinska, M., Ciamporova, M. & Ovecka, M. 2012 Early Zn^2+^-induced effects on membrane potential account for primary heavy metal susceptibility in tolerant and sensitive Arabidopsis species. Annals of botany 110, 445-459. (DOI:10.1093/aob/mcs111).

19. Nadgorska-Socha, A., Ptasinski, B. & Kita, A. 2013 Heavy metal bioaccumulation and antioxidative responses in *Cardaminopsis arenosa* and *Plantago lanceolata* leaves from metalliferous and non-metalliferous sites: a field study. Ecotoxicology 22, 1422-1434. (DOI:10.1007/s10646-013-1129-y).

20. Hanikenne, M. & Nouet, C. 2011 Metal hyperaccumulation and hypertolerance: a model for plant evolutionary genomics. Curr Opin Plant Biol 14, 252-259.

21. Verbruggen, N., Hermans, C. & Schat, H. 2009 Molecular mechanisms of metal hyperaccumulation in plants. New Phytol 181, 759-776. (DOI:10.1111/j.1469-8137.2008.02748.x).

22. Douglas, B., Martin, M., Ben, B. & Steve, W. 2015 Fitting Linear Mixed-Effects Models Using lme4. Journal of Statistical Software 67, 1-48. (DOI:doi:10.18637/jss.v067.i01.).

23. John, F., Sanford, W. & 2011 An {R} Companion to Applied Regression. Second Edition ed. Thousand Oaks, CA, Sage.

24. Rogstad, S. H. 1992 Saturated NaCl-CTAB Solution as a Means of Field Preservation of Leaves for DNA Analyses. Taxon 41, 701-708. (DOI:10.2307/1222395).

25. McKenna, A., Hanna, M., Banks, E., Sivachenko, A., Cibulskis, K., Kernytsky, A., Garimella, K., Altshuler, D., Gabriel, S., Daly, M., et al. 2010 The Genome Analysis Toolkit: a MapReduce framework for analyzing next-generation DNA sequencing data. Genome Res 20, 1297-1303. (DOI:10.1101/gr.107524.110).

26. Lê, S., Josse, J. & Husson, F. 2008 FactoMineR: An R Package for Multivariate Analysis. 2008 25, 18. (DOI:10.18637/jss.v025.i01).

27. Yant, L., Hollister, J. D., Wright, K. M., Arnold, B. J., Higgins, J. D., Franklin, F. C. H. & Bomblies, K. 2013 Meiotic adaptation to genome duplication in *Arabidopsis arenosa*. Curr Biol 23, 2151-2156. (DOI:10.1016/j.cub.2013.08.059).

28. Arnold, B. J., Lahner, B., DaCosta, J. M., Weisman, C. M., Hollister, J. D., Salt, D. E., Bomblies, K. & Yant, L. 2016 Borrowed alleles and convergence in serpentine adaptation. PNAS 113, 8320-8325. (DOI:www.pnas.org/cgi/doi/10.1073/pnas.1600405113).

29. Ross-Ibarra, J., Wright, S. I., Foxe, J. P., Kawabe, A., DeRose-Wilson, L., Gos, G., Charlesworth, D. & Gaut, B. S. 2008 Patterns of polymorphism and demographic history in natural populations of *Arabidopsis lyrata*. PLoS One 3, e2411. (DOI:10.1371/journal.pone.0002411).

30. Wright, S. 1951 The genetical structure of populations. Ann Eugen 15, 323-354.

31. Nielsen, R., Hubisz, M. J., Hellmann, I., Torgerson, D., Andres, A. M., Albrechtsen, A., Gutenkunst, R., Adams, M. D., Cargill, M., Boyko, A., et al. 2009 Darwinian and demographic forces affecting human protein coding genes. Genome Res 19, 838-849. (DOI:10.1101/gr.088336.108).

32. Smith, J. & Kronforst, M. R. 2013 Do Heliconius butterfly species exchange mimicry alleles? Biol Lett 9, 20130503. (DOI:10.1098/rsbl.2013.0503).

33. Bonhomme, M., Chevalet, C., Servin, B., Boitard, S., Abdallah, J., Blott, S. & SanCristobal, M. 2010 Detecting Selection in Population Trees: The Lewontin and Krakauer Test Extended. Genetics 186, 241-262. (DOI:10.1534/genetics.110.117275).

34. Teo, Y. Y., Fry, A. E., Bhattacharya, K., Small, K. S., Kwiatkowski, D. P. & Clark, T. G. 2009 Genome-wide comparisons of variation in linkage disequilibrium. Genome Res 19, 1849-1860. (DOI:10.1101/gr.092189.109).

35. Tajima, F. 1989 Statistical Method for Testing the Neutral Mutation Hypothesis by DNA Polymorphism. Genetics 123, 585-595.

36. Fay, J. C. & Wu, C.-I. 2000 Hitchhiking Under Positive Darwinian Selection. Genetics 155, 1405-1413.

37. Pavlidis, P., Zivkovic, D., Stamatakis, A. & Alachiotis, N. 2013 SweeD: likelihood-based detection of selective sweeps in thousands of genomes. Mol Biol Evol 30, 2224-2234. (DOI:10.1093/molbev/mst112).

38. Cingolani, P., Platts, A., Wang le, L., Coon, M., Nguyen, T., Wang, L., Land, S. J., Lu, X. & Ruden, D. M. 2012 A program for annotating and predicting the effects of single nucleotide polymorphisms, SnpEff: SNPs in the genome of *Drosophila melanogaster* strain w1118; iso-2; iso-3. Fly (Austin) 6, 80-92. (DOI:10.4161/fly.19695).

39. Rawat, V., Abdelsamad, A., Pietzenuk, B., Seymour, D. K., Koenig, D., Weigel, D., Pecinka, A. & Schneeberger, K. 2015 Improving the Annotation of *Arabidopsis lyrata* Using RNA-Seq Data. PLoS One 10, e0137391. (DOI:10.1371/journal.pone.0137391).

40. Oliveros, J. C. 2007-2015 Venny. An interactive tool for comparing lists with Venn’s diagrams. http://bioinfogp.cnb.csic.es/tools/venny/index.html.

41. Bindea, G., Mlecnik, B., Hackl, H., Charoentong, P., Tosolini, M., Kirilovsky, A., Fridman, W. H., Pages, F., Trajanoski, Z. & Galon, J. 2009 ClueGO: a Cytoscape plug-in to decipher functionally grouped gene ontology and pathway annotation networks. Bioinformatics 25, 1091-1093. (DOI:10.1093/bioinformatics/btp101).

42. Shannon, P., Markiel, A., Ozier, O., Baliga, N. S., Wang, J. T., Ramage, D., Amin, N., Schwikowski, B. & Ideker, T. 2003 Cytoscape: A Software Environment for Integrated Models of Biomolecular Interaction Networks. Genome Research 13, 2498-2504.

43. Gunther, T. & Coop, G. 2013 Robust identification of local adaptation from allele frequencies. Genetics 195, 205-220. (DOI:10.1534/genetics.113.152462).

44. Jeffreys, H. 1961 The Theory of Probability. 3rd edn ed. Oxford, Oxford University Press.

45. Šali, A. & Blundell, T. L. 1993 Comparative Protein Modelling by Satisfaction of Spatial Restraints. Journal of Molecular Biology 234, 779-815. (DOI:https://doi.org/10.1006/jmbi.1993.1626).

46. Buchan, D. W. A., Minneci, F., Nugent, T. C. O., Bryson, K. & Jones, D. T. 2013 Scalable web services for the PSIPRED Protein Analysis Workbench. Nucleic Acids Research 41, W349-W357. (DOI:10.1093/nar/gkt381).

47. Waterhouse, A. M., Procter, J. B., Martin, D. M. A., Clamp, M. & Barton, G. J. 2009 Jalview Version 2—a multiple sequence alignment editor and analysis workbench. Bioinformatics 25, 1189-1191. (DOI:10.1093/bioinformatics/btp033).

48. Briskine, R. V., Paape, T., Shimizu-Inatsugi, R., Nishiyama, T., Akama, S., Sese, J. & Shimizu, K. K. 2017 Genome assembly and annotation of *Arabidopsis halleri,* a model for heavy metal hyperaccumulation and evolutionary ecology. Mol Ecol Resour 17, 1025-1036. (DOI: 10.1111/1755-0998.12604).

49. Pauwels, M., Frerot, H., Bonnin, I. & Saumitou-Laprade, P. 2006 A broad-scale analysis of population differentiation for Zn tolerance in an emerging model species for tolerance study: *Arabidopsis halleri* (Brassicaceae). J Evol Biol 19, 1838-1850. (DOI:10.1111/j.1420-9101.2006.01178.x).

50. Long, T. A., Tsukagoshi, H., Busch, W., Lahner, B., Salt, D. E. & Benfey, P. N. 2010 The bHLH transcription factor POPEYE regulates response to iron deficiency in Arabidopsis roots. Plant Cell 22, 2219-2236. (DOI:10.1105/tpc.110.074096).

51. Bernal, M., Casero, D., Singh, V., Wilson, G. T., Grande, A., Yang, H., Dodani, S. C., Pellegrini, M., Huijser, P., Connolly, E. L., et al. 2012 Transcriptome sequencing identifies SPL7-regulated copper acquisition genes FRO4/FRO5 and the copper dependence of iron homeostasis in Arabidopsis. Plant Cell 24, 738-761. (DOI:10.1105/tpc.111.090431).

52. Gayomba, S. R., Jung, H. I., Yan, J., Danku, J., Rutzke, M. A., Bernal, M., Krämer, U., Kochian, L. V., Salt, D. E. & Vatamaniuk, O. K. 2013 The CTR/COPT-dependent copper uptake and SPL7-dependent copper deficiency responses are required for basal cadmium tolerance in *A. thaliana*. Metallomics 5, 1262-1275. (DOI:10.1039/c3mt00111c).

53. Connolly, E. L., Fett, J. P. & Guerinot, M. L. 2002 Expression of the IRT1 metal transporter is controlled by metals at the levels of transcript and protein accumulation. Plant Cell 14, 1347-1357.

54. Vert, G., Grotz, N., Dedaldechamp, F., Gaymard, F., Guerinot, M. L., Briat, J. F. & Curie, C. 2002 IRT1, an Arabidopsis transporter essential for iron uptake from the soil and for plant growth. Plant Cell 14, 1223-1233.

55. Arrivault, S., Senger, T. & Krämer, U. 2006 The Arabidopsis metal tolerance protein AtMTP3 maintains metal homeostasis by mediating Zn exclusion from the shoot under Fe deficiency and Zn oversupply. Plant J 46, 861-879. (DOI:10.1111/j.1365-313X.2006.02746.x).

56. Connolly, E. L., Campbell, N. H., Grotz, N., Prichard, C. L. & Guerinot, M. L. 2003 Overexpression of the FRO2 ferric chelate reductase confers tolerance to growth on low iron and uncovers posttranscriptional control. Plant Physiol 133, 1102-1110. (DOI:10.1104/pp.103.025122).

57. Cosio, C., DeSantis, L., Frey, B., Diallo, S. & Keller, C. 2005 Distribution of cadmium in leaves of *Thlaspi caerulescens*. Journal of experimental botany 56, 765-775. (DOI:10.1093/jxb/eri062).

58. Kovalchuk, O., Titov, V., Hohn, B. & Kovalchuk, I. 2001 A sensitive transgenic plant system to detect toxic inorganic compounds in the environment. Nat Biotechnol 19, 568-572. (DOI:10.1038/89327).

59. Hanikenne, M., Talke, I. N., Haydon, M. J., Lanz, C., Nolte, A., Motte, P., Kroymann, J., Weigel, D. & Krämer, U. 2008 Evolution of metal hyperaccumulation required *cis-*regulatory changes and triplication of *HMA4*. Nature 453, 391-395.

60. Dräger, D. B., Desbrosses-Fonrouge, A. G., Krach, C., Chardonnens, A. N., Meyer, R. C., Saumitou-Laprade, P. & Krämer, U. 2004 Two genes encoding *Arabidopsis halleri* MTP1 metal transport proteins co-segregate with zinc tolerance and account for high MTP1 transcript levels. Plant J 39, 425-439.

61. Suryawanshi, V., Talke, I. N., Weber, M., Eils, R., Brors, B., Clemens, S. & Kramer, U. 2016 Between-species differences in gene copy number are enriched among functions critical for adaptive evolution in *Arabidopsis halleri*. BMC Genomics 17, 1034. (DOI:10.1186/s12864-016-3319-5).

62. Hanikenne, M., Kroymann, J., Trampczynska, A., Bernal, M., Motte, P., Clemens, S. & Krämer, U. 2013 Hard selective sweep and ectopic gene conversion in a gene cluster affording environmental adaptation. PLoS genetics 9, e1003707. (DOI:10.1371/journal.pgen.1003707).

63. Koizumi, M., Yamaguchi-Shinozaki, K., Tsuji, H. & Shinozaki, K. 1993 Structure and expression of two genes that encode distinct drought-inducible cysteine proteinases in *Arabidopsis thaliana*. Gene 129, 175-182.

64. Bernoux, M., Timmers, T., Jauneau, A., Briere, C., de Wit, P. J., Marco, Y. & Deslandes, L. 2008 RD19, an Arabidopsis cysteine protease required for RRS1-R-mediated resistance, is relocalized to the nucleus by the *Ralstonia solanacearum* PopP2 effector. Plant Cell 20, 2252-2264. (DOI:10.1105/tpc.108.058685).

65. Hamel, F., Breton, C. & Houde, M. 1998 Isolation and characterization of wheat aluminum-regulated genes: possible involvement of aluminum as a pathogenesis response elicitor. Planta 205, 531-538.

66. Usui, M., Tanaka, S., Miyasaka, H., Suzuki, Y. & Shioi, Y. 2007 Characterization of cysteine protease induced by oxidative stress in cells of *Chlamydomonas* sp. strain W80. Physiol. Plant. 131, 519-526.

67. Becher, M., Talke, I. N., Krall, L. & Krämer, U. 2004 Cross-species microarray transcript profiling reveals high constitutive expression of metal homeostasis genes in shoots of the zinc hyperaccumulator *Arabidopsis halleri*. The Plant journal: for cell and molecular biology 37, 251-268.

68. Woolhouse, H. W. 1983 Toxicity and Tolerance in the Response of Plants to Metals. In Encyclopedia of Plant Physiology, New Series (eds. L. O.L., P. S. Nobel, O. C.B. & H. Ziegler), pp. 245-300. Berlin, Germany, Springer-Verlag.

